# Human Missense Variation is Constrained by Domain Structure and Highlights Functional and Pathogenic Residues

**DOI:** 10.1101/127050

**Authors:** Stuart A. MacGowan, Fábio Madeira, Thiago Britto-Borges, Melanie S. Schmittner, Christian Cole, Geoffrey J. Barton

## Abstract

Human genome sequencing has generated population variant datasets containing millions of variants from hundreds of thousands of individuals^1-3^. The datasets show the genomic distribution of genetic variation to be influenced on genic and sub-genic scales by gene essentiality,^1,4,5^ protein domain architecture^6^ and the presence of genomic features such as splice donor/acceptor sites.^2^ However, the variant data are still too sparse to provide a comparative picture of genetic variation between individual protein residues in the proteome.^1,6^ Here, we overcome this sparsity for ∼25,000 human protein domains in 1,291 domain families by aggregating variants over equivalent positions (columns) in multiple sequence alignments of sequence-similar (paralagous) domains^7,8^. We then compare the resulting variation profiles from the human population to residue conservation across all species^9^ and find that the same tertiary structural and functional pressures that affect amino acid conservation during domain evolution constrain missense variant distributions. Thus, depletion of missense variants at a position implies that it is structurally or functionally important. We find such positions are enriched in known disease-associated variants (OR = 2.83, *p* ≈ 0) while positions that are both missense depleted and evolutionary conserved are further enriched in disease-associated variants (OR = 1.85, *p* = 3.3×10^-17^) compared to those that are only evolutionary conserved (OR = 1.29, *p* = 4.5×10^-19^). Unexpectedly, a subset of evolutionary Unconserved positions are Missense Depleted in human (UMD positions) and these are also enriched in pathogenic variants (OR = 1.74, *p* = 0.02). UMD positions are further differentiated from other unconserved residues in that they are enriched in ligand, DNA and protein binding interactions (OR = 1.59, *p* = 0.003), which suggests this stratification can identify functionally important positions. A different class of positions that are Conserved and Missense Enriched (CME) show an enrichment of ClinVar risk factor variants (OR = 2.27, *p* = 0.004). We illustrate these principles with the G-Protein Coupled Receptor (GPCR) family, Nuclear Receptor Ligand Binding Domain family and In Between Ring-Finger (IBR) domains and list a total of 343 UMD positions in 211 domain families. This study will have broad applications to: (a) providing focus for functional studies of specific proteins by mutagenesis; (b) refining pathogenicity prediction models; (c) highlighting which residue interactions to target when refining the specificity of small-molecule drugs.

## Variant densities and the sparsity problem

Human sequencing projects are beginning to shed light on the patterns of genetic variation that are present in human populations.^1,2^ One way in which these studies enhance the understanding of inter-individual variation is by characterising different densities of single-nucleotide variants (SNVs) and short insertion and deletions (indels) at different genomic loci. Analysis of large cohort variation datasets has revealed that genes differ in their tolerance of non-synonymous and loss-of-function variation.^1,4^ Within protein-coding genes, regions that encode protein domains are less tolerant of non-synonymous variants than inter-domain coding regions and are more prone to disease variants.^6^ The 60,706 sample Exome Aggregation Consortium1 study yielded ∼125 variants per kilobase, rendering a per nucleotide comparison impossible since most single nucleotides have zero variants. Variant sparsity can also be addressed by aggregating over pseudo-paralogous positions. For example, aligning nucleotide sequences on start codons reveals that start codons have fewer variants than adjacent sites, while the 5’-UTR is more variable than the CDS and every third base in a codon variable.2 These differences are observed because the pressures imposed by those genomic features are common to each individual aligned sequence.

## Residue resolution through protein family aggregation

Multiple sequence alignments (MSA) are a well established way to identify position-specific features in a family of homologous sequences. Figure 1A illustrates schematically how an MSA containing multiple human paralogs can be used to aggregate SNVs from multiple loci in a position specific manner. This process condenses the sparse variant counts from single sequences into dense variant counts for the domain family. Similar approaches have been adopted to identify low frequency cancer driver mutations,^10-12^ and find sites in domains where pathogenic mutations cluster.^13^ To perform a comprehensive analysis of protein domains, germline variation data retrieved from Ensembl^14,15^ was aggregated with respect to the domain families in Pfam.^8^ Pfam contains 16,035 domain families and of these families 6,088 contain at least one human sequence and 1,376 have at least five after adjusting for duplicate sequences (see Methods). Figures 1B-C show that even though most human sequence residues in Pfam domains have zero variants, after aggregation most Pfam domain family positions have at least two variants.

**Figure 1:**
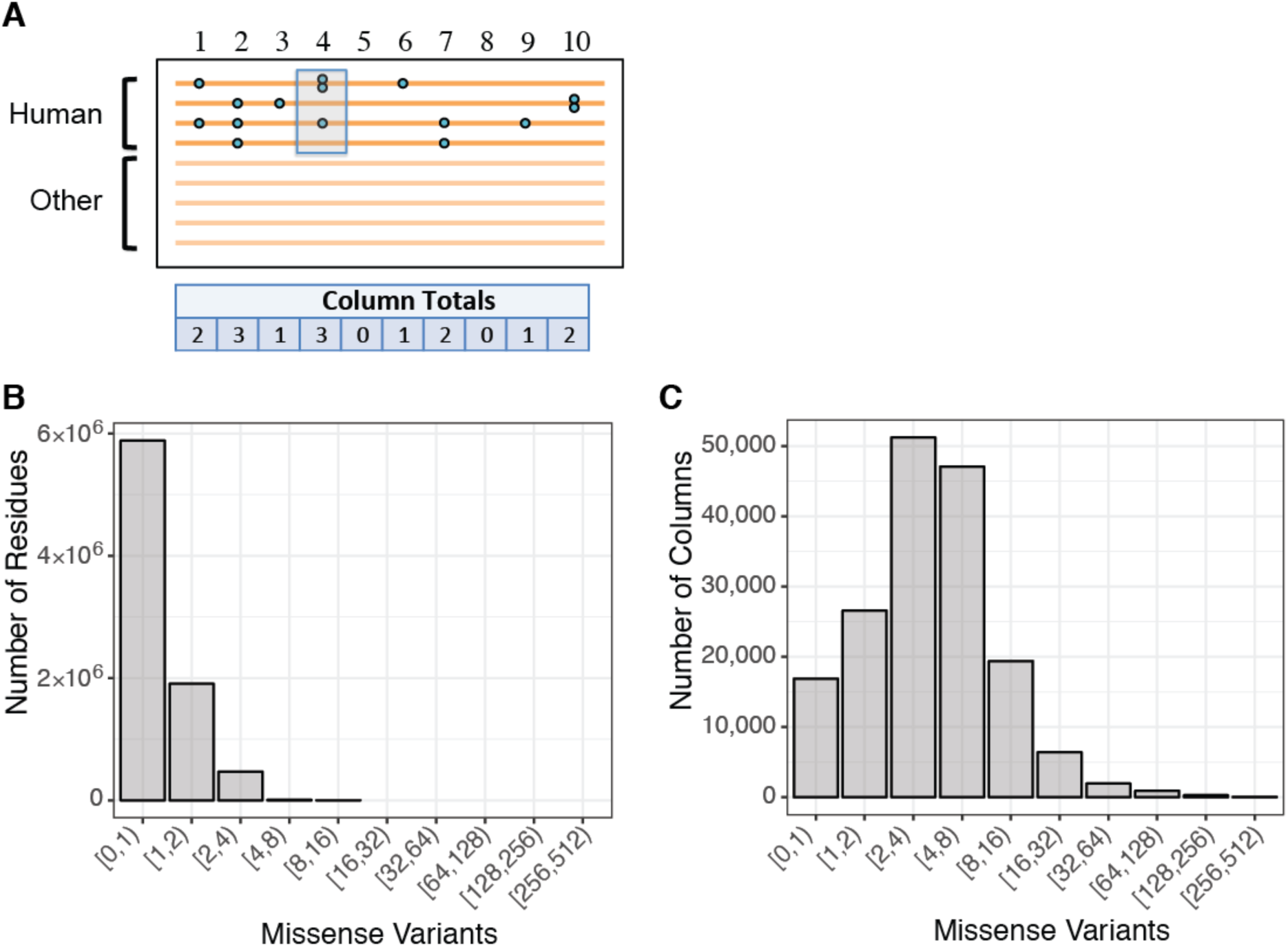
Variant aggregation over protein family alignments. A. Schematic illustration of a protein family alignment. Each line represents a human or non-human sequence and human sequences can have zero or more variants (blue circles). Few variants are observed at each alignment position per sequence but the column totals are larger. B. Distribution of variants per human residue in all Pfam sequences (2,927,499 missense variants, 8,264,091 residues; no filters applied). C. Distribution of variants per alignment column in Pfam alignments (955,636 missense variants, 159,296 columns; includes only columns with at least five human residues).

## SNV density is correlated with evolutionary conservation

Accurate predictions of structure and function can be made from MSAs^16-18^ because these features impose constraints on accepted mutations in domain families. These constraints can be inferred from patterns in residue conservation scores,^9^ which quantify the extent of residue or physicochemical property conservation at each position in the alignment. In protein domain family MSAs, which can contain orthologs and paralogs in varying proportions, these scores are interpreted as the degree of evolutionary conservation in each site of the domain family and are different to conservation scores for alignments that contain only closely related orthologs because of greater functional divergence. Throughout this text, the term evolutionary conservation refers to the conservation of residues during domain family evolution and accounts for orthologous and paralogous evolutionary process as captured in the Pfam alignments.

Figure 2A shows the correlation between the domain family column variant counts and the Shenkin divergence score (V_Shenkin_)^19^ in the SH2 domain family (PF00017). The number of missense variants increases with increasing residue divergence (i.e., decreasing conservation) whilst the frequency of synonymous variation remains constant with respect to column conservation. Extended Data Figs. 1 and 2 illustrate this behaviour on the SH2 alignment and crystal structure and show that in this example, the protein’s secondary and tertiary structures and domain-domain interactions are common factors constraining both conservation and population constraint. This demonstrates that the missense variant distribution is subject to the same structural and functional constraints over generational timescales that affect amino acid substitution frequencies over evolutionary timescales. In contrast, the distribution of synonymous variation is not affected because these variants are silent at the protein structure level. Figure 2B shows that this result extends to other protein families by illustrating that the V_Shenkin_ regression coefficients for each family are distributed around zero for synonymous variant totals and are typically positive for missense variants.

**Figure 2:**
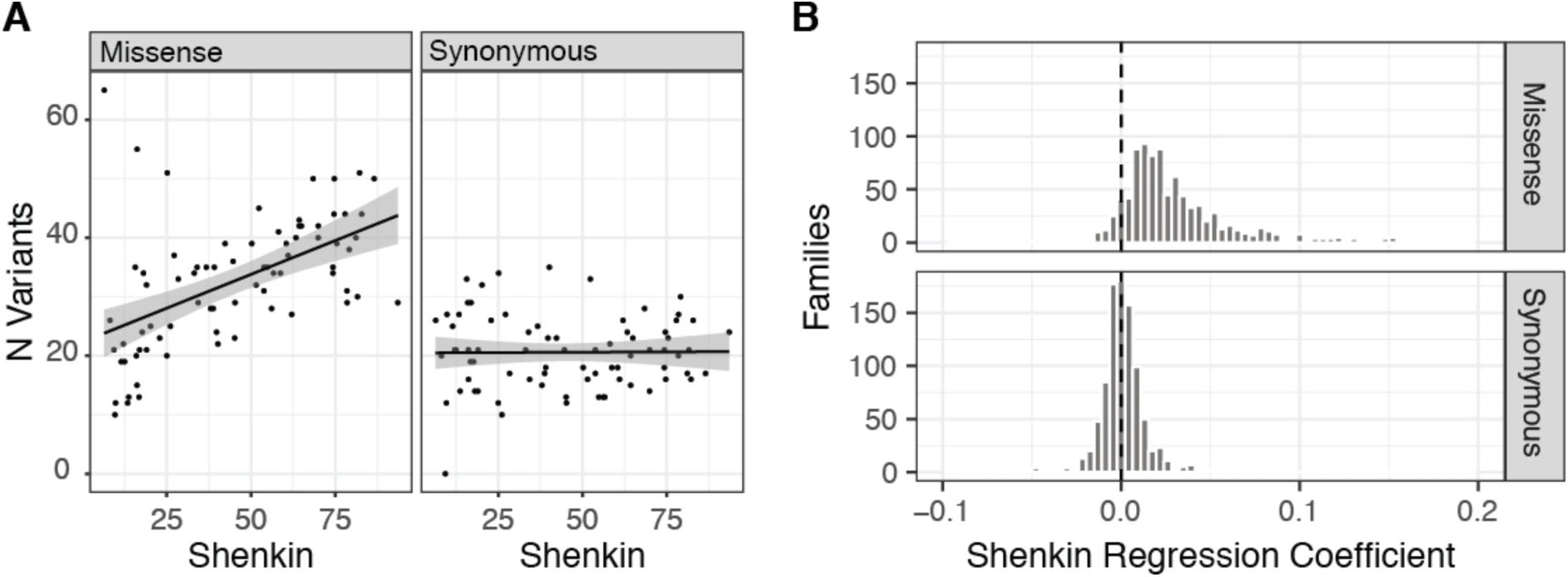
Relationship between column variant totals and VShenkin. A. Variant counts vs. VShenkin for missense (left panel) and synonymous variants (right panel) for the SH2 domain (PF00017). The regression lines show least-squares fits and the shaded regions indicate standard errors of prediction. B. Histograms showing the distributions of VShenkin regression coefficients for linear models fitting column variant totals to VShenkin and column human residue occupancies for protein families with > 50 included alignment columns (n = 934).

## Properties of sites relatively depleted or enriched for missense variation

Domain family alignment columns were classified as missense depleted or missense enriched by testing whether a column possessed significantly more or less missense variation than observed elsewhere in the alignment (see Methods). Figure 3A shows that with respect to ClinVar^20^ variant annotations missense depleted columns have higher rates of ‘pathogenic’ (Fisher OR = 2.83, *p* ≈ 0) and ‘likely pathogenic’ variants (OR = 2.17, *p* = 1.9×10^-12^) compared to other sites, indicating that diversity is suppressed in positions that are critical for function. Variant enriched columns possess proportionally more ‘risk factor’ variants (Fisher OR = 1.66, *p* = 0.017). This may suggest that there is generally an increased chance of co-segregating phenotypic differences at sites with relatively high population diversity.

**Figure 3:**
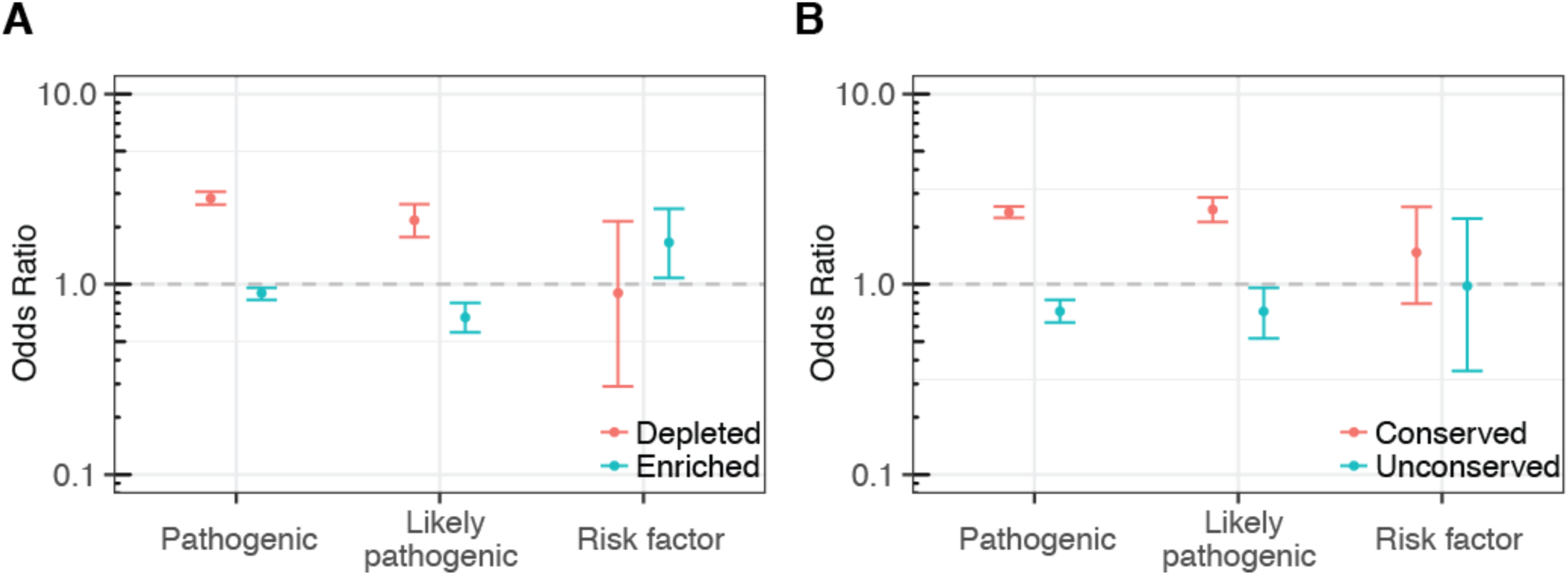
Properties of missense depleted and enriched domain family alignment columns. Odds ratios and 95% C.I. for enrichment of variants with specific ClinVar terms that affect residues found in A. missense depleted (*p* < 0.1; see methods) or enriched (*p* < 0.1) domain family alignment columns and B. conserved (C_Valdar_ in 1^st^ decile) or unconserved columns (C_Valdar_ in 10^th^ decile).

For comparison, Figure 3B shows the equivalent ClinVar association tests for columns classified by their evolutionary conservation as measured by Valdar’s score (C_Valdar_).^9^ For pathogenic variants, conserved vs. unconserved columns display the same behaviour as missense depleted vs. enriched columns, which is concordant with previous work and expected since most missense depleted columns are also conserved. However, the column classification schemes yield almost opposite trends with respect to the distribution of ClinVar risk factor variants. There is a slight tendency for risk factor variants to occur more frequently in evolutionary conserved columns (OR = 1.47, *p* = 0.194), which contrasts with their higher frequencies in columns that are relatively enriched for missense variation.

## The Conservation Plane: Combining column variant class and conservation

Although the distribution of missense variants within domains is typically concordant with the evolutionary conservation profile (Figure 2B), the two metrics are not redundant and cross-classification of alignment columns by both yields residue categories with interesting properties. Figure 4A shows the distribution of ClinVar annotated pathogenic variants between columns classified as unconserved-missense depleted (UMD), unconserved-missense enriched (UME), conserved-missense depleted (CMD) and conserved-missense enriched (CME). Conserved and unconserved columns that are neither missense depleted or enriched, i.e. have an average number of missense variants for the family, are also shown. It shows that: 1) all conserved sites are enriched for pathogenic variants but CMD sites are more so (CME: OR = 1.24, *p* = 1.6×10^-5^; CMD: OR = 1.85, *p* = 3.3×10^-17^) and 2) the UMD subset of unconserved residues are enriched for pathogenic variants to an extent comparable to conserved residues (OR = 1.74, *p* = 0.02). The UMD classification identifies sites where residues have varied throughout the evolution of the domain family but the specific residue adopted by each domain is now under negative selection in human. This implies that residues in this column class could be enriched for specificity determinants. A structural analysis of 270 UMD sites found in 160 families provides some support for this hypothesis. We compared these sites to UME columns from the same families and found that UMD columns were enriched for ligand, domain-domain and nucleotide interactions (OR = 1.59, *p* = 0.003) and tended to be less accessible to solvent (OR = 1.73, *p* = 2.0 × 10^-04^; Extended Data Table 1). Figure 4C illustrates an example of a protein family where UMD residues indicate known ligand-binding sites. The Rhodopsin-like receptor family (PF00001) contains 11 UMD sites, five of which occur in sequence in the centre of Helix 3 and form interactions with ligands in many structures (e.g. residues in column 780 interact with ligands in 23 distinct proteins; Extended Data Table 2) and includes a Na^2+^ binding residue. Extended Data Fig. 3 shows another example of ligand binding site identification in the nuclear receptor ligand binding domain family (NR-LBD; PF00104). Additionally, Extended Data Fig. 4 shows UMD sites in the NR-LBD family that are not directly involved in ligand binding but instead mediate strong intra-domain cross-helical interactions that vary dramatically between domains. Structures of intact DNA-bound nuclear receptors suggest that in some proteins these residues interact with the LBD-DNA binding domain linker and thus may mediate the ligand induced DNA binding response (Not shown. For an example see Glu 295 and Ser 332 in PDB ID: 3e00 chain D.).^21^ These important interactions may not be detected by residue co-variation analysis^18^ because the UMD site interacts with residues aligned in different columns in each domain. One UMD site is seen in the IBR domain (PF01485). In the E3 ubiquitin-protein ligase parkin, this is Glu370 that recent structural studies suggest is at the interface with Ubiquitin^22^ and so likely to be important in mediating this interaction. All other UMD classified sites can be found in Supplementary Data Table 1. Together, these findings show that human missense variation can stratify unconserved alignment columns to identify a small number of residues likely to be important for function and specificity.

**Figure 4:**
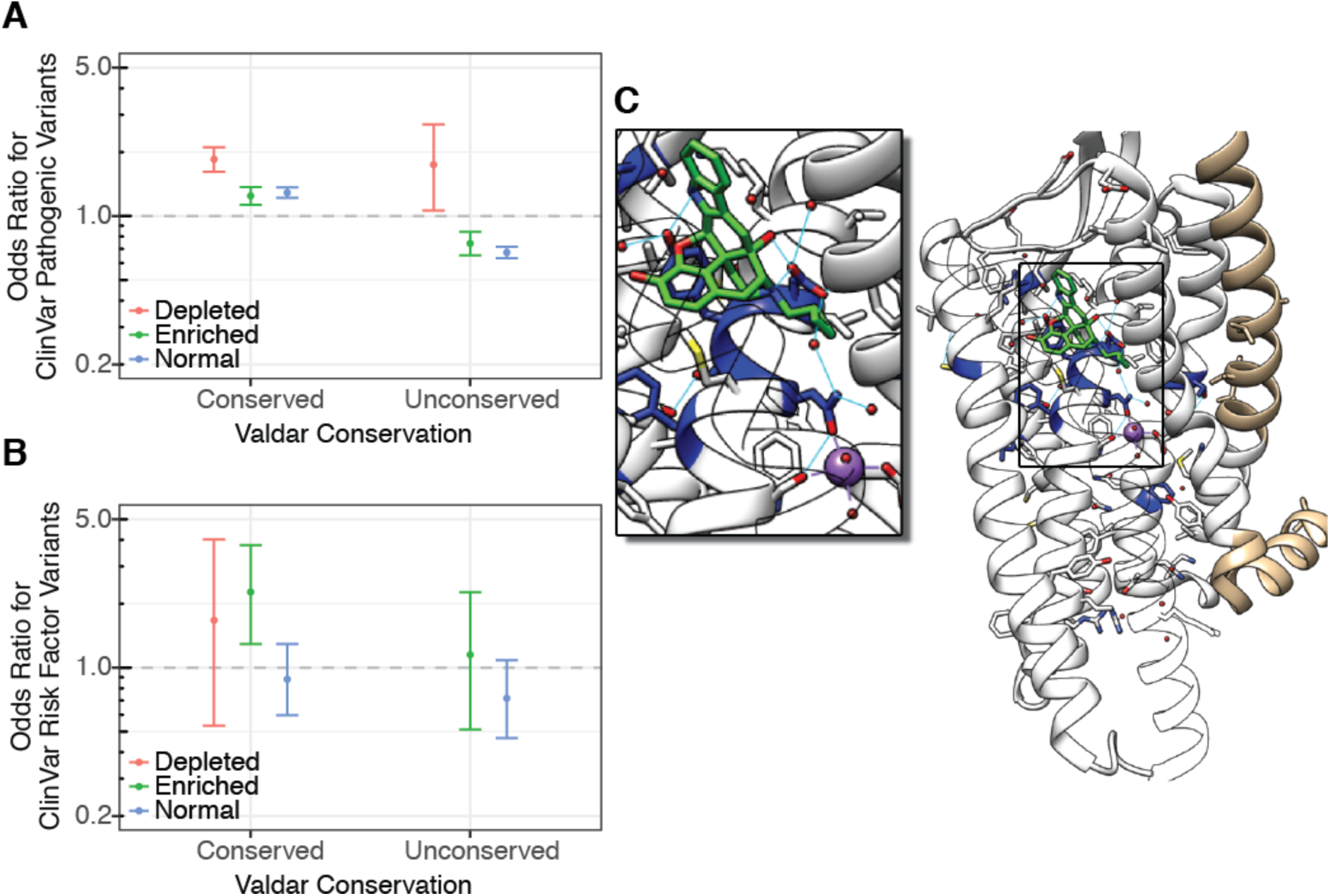
Classification of domain residues by evolutionary conservation and relative population variation. A. Odds ratios for ClinVar pathogenic variants in missense depleted (*p* < 0.1; see methods), enriched (*p* < 0.1) or normal (*p* ≥ 0.1) alignment columns that were either conserved (C_Valdar_ < median) or unconserved (C_Valdar_ > median). B. Odds ratios for ClinVar risk factor variants in different column classes. UMD columns are not shown as there are zero risk factor variants in this column class; the ClinVar risk factor OR and 95 % C.I. for UMD columns is 0 [0, 14]. C. Illustration of UMD residues (blue) in the Rhodopsin-like receptors (PF00001) mapped to a structure of the Delta-type opioid receptor (PDB ID: 4n6h).^23^ Amongst the 11 UMD residues are several involved in ligand binding and one that coordinates the bound sodium ion; residues 249-287 are hidden for clarity.

Another striking feature of residues in columns with discordant levels of evolutionary conservation and population diversity was found. Figure 4B shows the odds ratios of observing ClinVar risk factor variants in columns classed according to evolutionary conservation and whether they are relatively enriched in missense variants or not and highlights that CME sites are significantly enriched in risk factor variants (OR = 2.27, *p* = 0.004). This is consistent with the previous observation that missense enriched columns were enriched for risk factor variants and that conserved columns showed a tendency toward risk factor enrichment (Figure 3) but the combined effect is much stronger. To our knowledge this is the first time that a feature marking residues pre-disposed to carrying risk factor variants has been identified.

With further development, the conservation plane may yield insight into the evolutionary forces acting on individual sites in protein domain families. Although this will require consideration of each family’s phylogeny coupled with more detailed variation metrics (e.g., considering allele frequencies, heterozygosity, missense/synonymous ratios (d_N_/d_S_), McDonald–Kreitman test^24^ and derivatives) our results offer clues as to which evolutionary signatures are being detected. Given the recognised effects of different types of selection upon intra- and interspecific variability,^25^ we can loosely associate: CMD sites with negative selection and sites affected by selective sweeps; UMD sites with positive selection (here, domain specialisation) and CME sites with balancing selection. Whilst these associations are speculative, the structural features and disease associations of those classes are congruent with these evolutionary processes.^25-27^ A few immediate practical applications follow from the missense-depletion and conservation plane class associations. For variant pathogenicity prediction, the results extend the work of Gussow and coworkers^4,6^ and open the door to hierarchical classification where the impact of a variant can be can be judged in genic, sub-genic architecture, and now, residue level contexts on the basis of population variation. In protein feature prediction, the ability to identify functionally important residues that are classically unconserved could help to identify allosteric and surface interaction sites, whilst a metric that is sensitive to specificity determining residues should prove useful in understanding enzyme active sites and other functional sites in more detail.

## Methods

### Datasets, Mapping and Filtering

Protein family alignments were downloaded from Pfam (v29)^7,8^ and parsed using Biopython (v1.66, with patches #768 #769)^28^ and conservation scores were calculated by AACons via JABAWS (v2.1).^29^ The human sequences in the alignment were mapped to the corresponding full UniProt sequences to create keys between UniProt sequence residue numbers and Pfam alignment column numbers. For each human sequence, germline population variants were retrieved from Ensembl 84^14,15^ via the Ensembl API using ProteoFAV.^30^ Ensembl variants are provided with indexes to UniProt sequence residue numbers and were thus mapped to Pfam alignment columns.

Ensembl variation agglomerates variants and annotation data from a variety of sources including dbSNP (v146), 1KG, ESP and ExAC1. A full description of the variant sources present in Ensembl 84 is available at http://mar2016.archive.Ensembl.org/info/genome/variation/sources_documentation.html. Ensembl provides numerous annotations including the predicted protein consequences (i.e. missense, synonymous, stop gained, etc.), minor allele frequency (MAF) and ClinVar^20^ disease status. These annotations were used to filter the Pfam-mapped variants for the collection of variant sub-class alignment column statistics. For example, this is how the number of ClinVar ‘pathogenic’ missense variants in each alignment column was calculated.

Pfam (v29) contains 16,035 domain family alignments. Variants were gathered and mapped to the alignments for the 6,088 families that contain at least one human sequence. For inclusion in this analysis, a minimum threshold of five human sequences was adopted corresponding to 2,939 protein families. However, some of these families do not meet this criterion after sequence duplication correction (see below) leaving 1,376 families. Finally, alignment column conservation scores could not be obtained for 85 of the families, resulting in a final dataset of 1,291 protein families. These families contain an estimated 25,158 human protein domains. Only columns with ≥ 5 human residues (i.e., non-gap) were considered, corresponding to 159,296 alignment columns. This filter was applied in all analyses reported in this work.

### Variant Duplication

Some alignments contained variants that mapped to multiple sequences due to sequence duplication. For example, in PF00001 all variants that mapped to the human sequence P2Y11/45-321 (P2Y purinoceptor 11) from the P2RY11 gene are duplicated in A0A0B4J1V8/465-741 because this sequence contains the same 7 transmembrane receptor domain as P2Y11 as a result of A0A0B4J1V8 being the product of a read-through transcript that includes the P2RY11 gene. This means there are two copies of the P2RY11 7 transmembrane receptor domain in the alignment and its variant profile is doubly weighted. Another example in this family comes from human sequences MSHR/55-298 (Melanocortin receptor 1), G3V4F0/55-298 and A0A0B4J269, which all are mapped to the same genomic loci. Accordingly, sequence duplication was accounted for by de-duplicating variants and sequences before summing over columns.

### Statistical Analyses

The statistical analyses were all performed using R version 3.2.2. Regressions were calculated by the *lm* function from the *stats* library. Odds ratios and Fishers exact *p* values were calculated with the *fisher.test* function from the *stats* library. Plots were produced with *ggplot2*.

### Alignment Column Classification

Columns were classified as depleted, enriched or neutral with respect to the column variant totals relative to the average for the other columns in the alignment. For each alignment column *x*, a 2×2 table was constructed of the form *a, b, c, d* with elements: *a*. the number of variants mapped to residues in column *x, b*. the total number of variants mapped to all other alignment columns, *c*. the number of human residues in column *x* and *d*. the total number of human residues in the rest of the alignment. Application of the R *stats* function *fisher.test* to each table yielded an odds ratio > 1 if the column contained more than the alignment average number of variants per human residue or OR < 1 if there were fewer than the average number of variants per human residue. The function also provided the *p* value afforded by Fisher’s exact test. This meant that for a given *p*_*threshold*_ columns with *p* ≥ *p*_*threshold*_ were considered normal and columns with *p* < *p*_*threshold*_ were considered depleted if OR < 1 or enriched if OR > 1. Notably, in addition to the effect size, *p* is sensitive to data availability (i.e., variant counts) and alignment column occupancy. In this work, *p*_*threshold*_ = 0.1 unless otherwise specified.

### Structural Analysis of Evolutionary Unconserved and Missense Depleted Residues

Columns were classified as unconserved-missense depleted (UMD) or unconserved-missense enriched (UME) if they displayed significant residue diversity (V_Shenkin_ in 4^th^ quartile) and were missense depleted or enriched, respectively. The 343 columns in 211 families that met these criteria were subjected to an automated analysis where the flagged residues were mapped to PDB structures via SIFTS;^31^ 270 columns from 160 families were mapped to at least one PDB structure. Biological units were obtained from the PDBe in mmCIF format. When multiple biological units were available for a particular asymmetric unit, the preferred biological unit ID was obtained by querying the PDBe API.^32^ Atoms were considered to interact if they were within 5 Å. A residue was considered to participate in a domain interaction if it interacted with a Pfam domain on a different PDB chain. Residue relative solvent accessibilities (RSAs) were calculated from the DSSP accessible surface^33^ as described in Tien *et al*.^34^ and were classified as surface (RSA > 25%), partially exposed (5% < RSA ≤ 25%) or core (RSA ≤ 5%).

The results of the automated analysis were supplemented by a manual structural analysis using a workflow enabled by the Jalview multiple sequence alignment workbench^35^ and the UCSF Chimera molecular graphics program.^36^ Jalview feature files identifying the UMD columns were generated. When the feature files were loaded onto the appropriate alignment in Jalview, the residues in the UMD columns were highlighted for the user. Jalview was then used to find PDB structures for the sequences in the alignment that were then visualised in UCSF Chimera. Jalview automatically mapped the UMD residue annotations to the PDB structure so that the residues could be assessed in their structural context. UCSF Chimera was used to identify other residues in the structure that were hydrogen bonded to, or had a Van der Waals distance < 1 Å with, a side-chain atom of any UMD residues present. The residues were then classified according to any contacts made as either: ligand binding, ion binding, inter-domain interaction, intra-domain interaction or surface residue. This analysis found that of those families with UMD residues, 19% had at least one UMD site involved in ligand-binding whilst 42% had a site directly involved in domain-domain interactions.

## Code availability

The code used in this study is available from the Barton Group GitHub repository at https://github.com/bartongroup/. The software was not designed for portability and may not function as intended in all environments, but the source code illustrates our methodology. We are currently developing a production version that will enable users to apply our methods to their own alignments to be released in the same repository.

## Data availability

The multiple sequence alignments and human variation data that underlie and support the findings of this study are available from Pfam, http://pfam.xfam.org/ and Ensembl 84, http://www.Ensembl.org/, respectively. The calculated data, including alignment column variation statistics and residue conservation scores are presently available from the corresponding author upon request whilst a web resource is under development. The UMD columns are also identified in the supplementary material.

## Acknowledgements

We thank the Jalview development team for their help with streamlining the visualisation of alignment and structural data and Jim Procter for additional assistance in using AACons and useful discussions regarding evolutionary theory in relation to multiple sequence alignments. We also thank Helen Walden, Maurice van Steensel, Ulrich Zachariae, Owen Vickery, David Gray and Alessio Ciulli for discussions about specific protein families. This work was supported by Wellcome Trust Strategic Awards [098439/Z/12/Z and WT097945], Wellcome Trust Doctoral Training Account [100150/Z/12/Z], Wellcome Trust Biomedical Resources Grant [101651/Z/13/Z], Coordenação de Aperfeiçoamento de Pessoal de Nível Superior studentship [CAPES process 1529/12-9] and Biotechnology and Biological Sciences Research Council Grants [BB/J019364/1, BB/L020742/1].

## Author contributions

S.A.M. designed and performed the study, analysed data and wrote the manuscript. F.M. contributed software to collect variation data and collected the interaction and RSA data for UMD and UME residues. T.B. contributed software to collect variation data. M.S. performed the manual structural analysis of UMD residues. C.C. analysed data. G.J.B. designed the study, analysed data and wrote the manuscript.

## Extended Data Captions

**Extended Data Figure 1:**
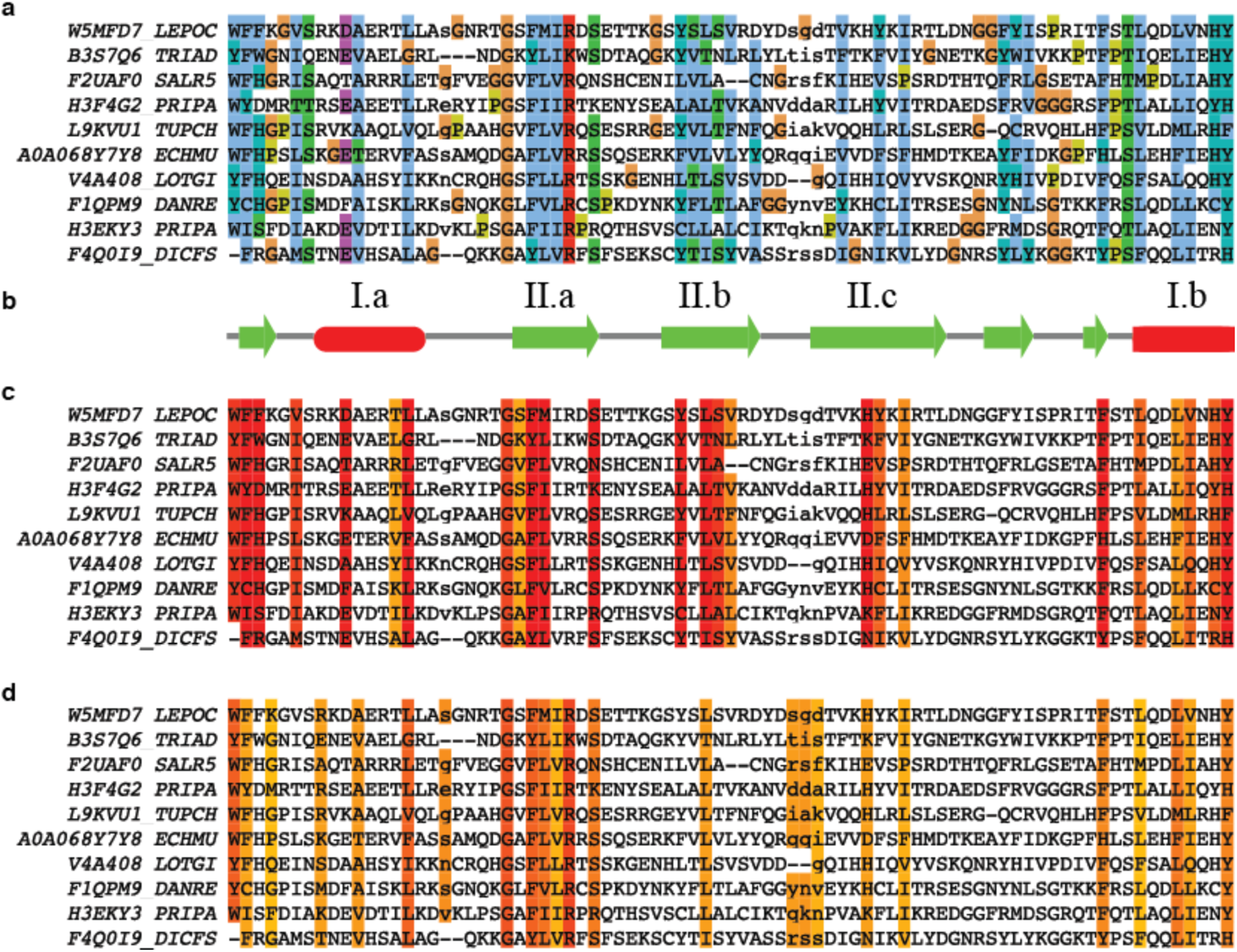
An extract of the SH2 alignment (PF00017.21) showing the influence of secondary structure constraints upon evolutionary conservation and missense depletion. a. Alignment with Clustal X^37^ colouring where blue indicates hydrophobic residue conservation. b. Consensus secondary structure from Pfam (v31);^7,8^ labelled elements indicate the archetypal SH2 partially buried helices (I.a and I.b) and β-strands (II.a-c). c. Missense depleted columns with *P* ≤ 0.2. d. Columns with V_Shenkin_ ≤ 20. The pattern of conserved hydrophobic residues in a are indicative of the structural constraints imposed by the secondary structure elements in b. These structural constraints are known to produce patterns in conservation metrics like V_Shenkin_ in d. These constraints also influence the distribution of missense depleted columns in c. Figure created with Jalview.^35^

**Extended Data Figure 2:**
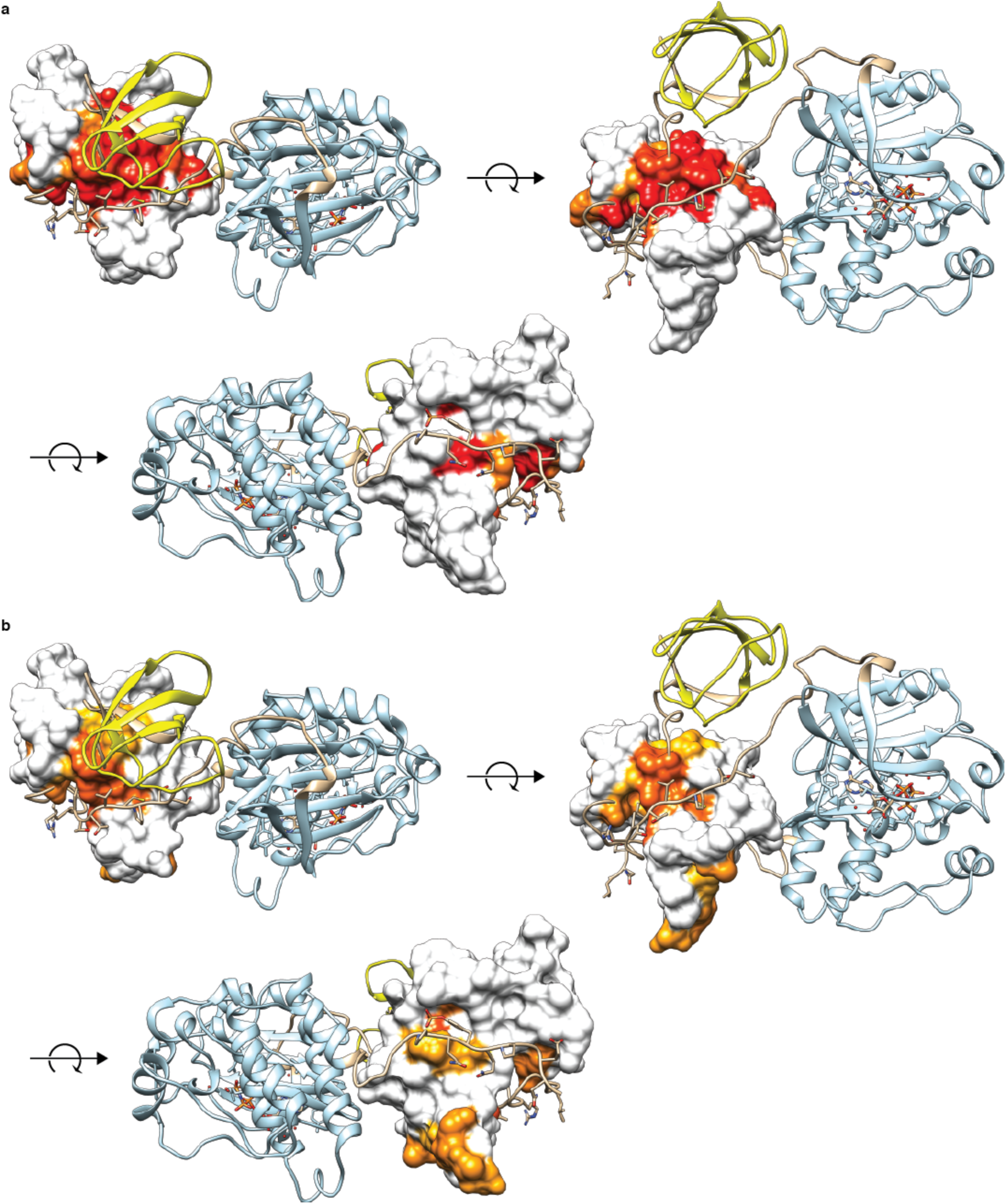
Inter-domain interactions of the SH2 domain in inactivated Src (PDB ID: 2src).^38^ The surface of the SH2 domain (PF00017) is coloured red to yellow corresponding to a. missense depletion *P* over range [0, 0.2) and b. V_Shenkin_ over range [0, 20); white surface regions are outside these ranges. The sub-panels show interactions with the Src SH3 domain (yellow), kinase-SH2 linker (tan) and the tail region including phosphorylated-Tyr (tan). Residues that interact with the SH2 domain are displayed as sticks. Figure created with Jalview^35^ and UCSF Chimera.^36^

**Extended Data Figure 3:**
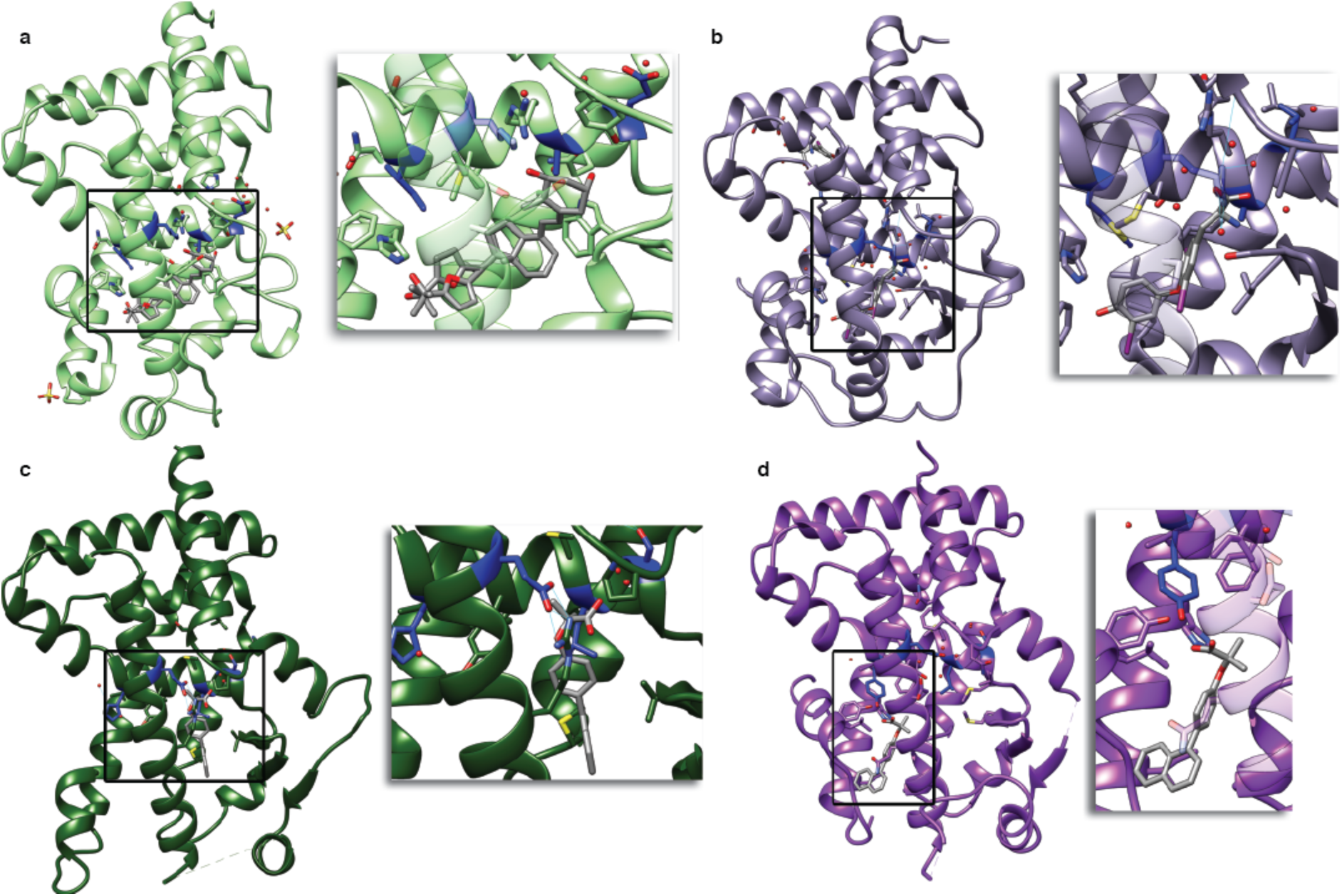
Examples of UMD residues (blue) involved in ligand-binding in the nuclear receptor ligand binding domains protein family (PF00104). a. VDR in complex with a calcitriol analog (3ogt).^39^ b. TH*α* in complex with triiodothyronine (4lnx).^40^ c. PPAR*γ* (5hzc) and d. PPAR*α* (5hyk) in complex with the PPAR pan-agonist AL29-26.^41^ The ligand is in VdW contact with the unconserved-depleted L330 in PPAR*γ* and with Y314 in PPAR*α*. Note that the substitution at the unconserved-depleted site H323 in PPAR*γ* to Y314 in PPAR*α* is related to the activity specificity of these two receptors with respect to AL29-26.^41^ Figure created with UCSF Chimera^36^ and Jalview.^35^

**Extended Data Figure 4:**
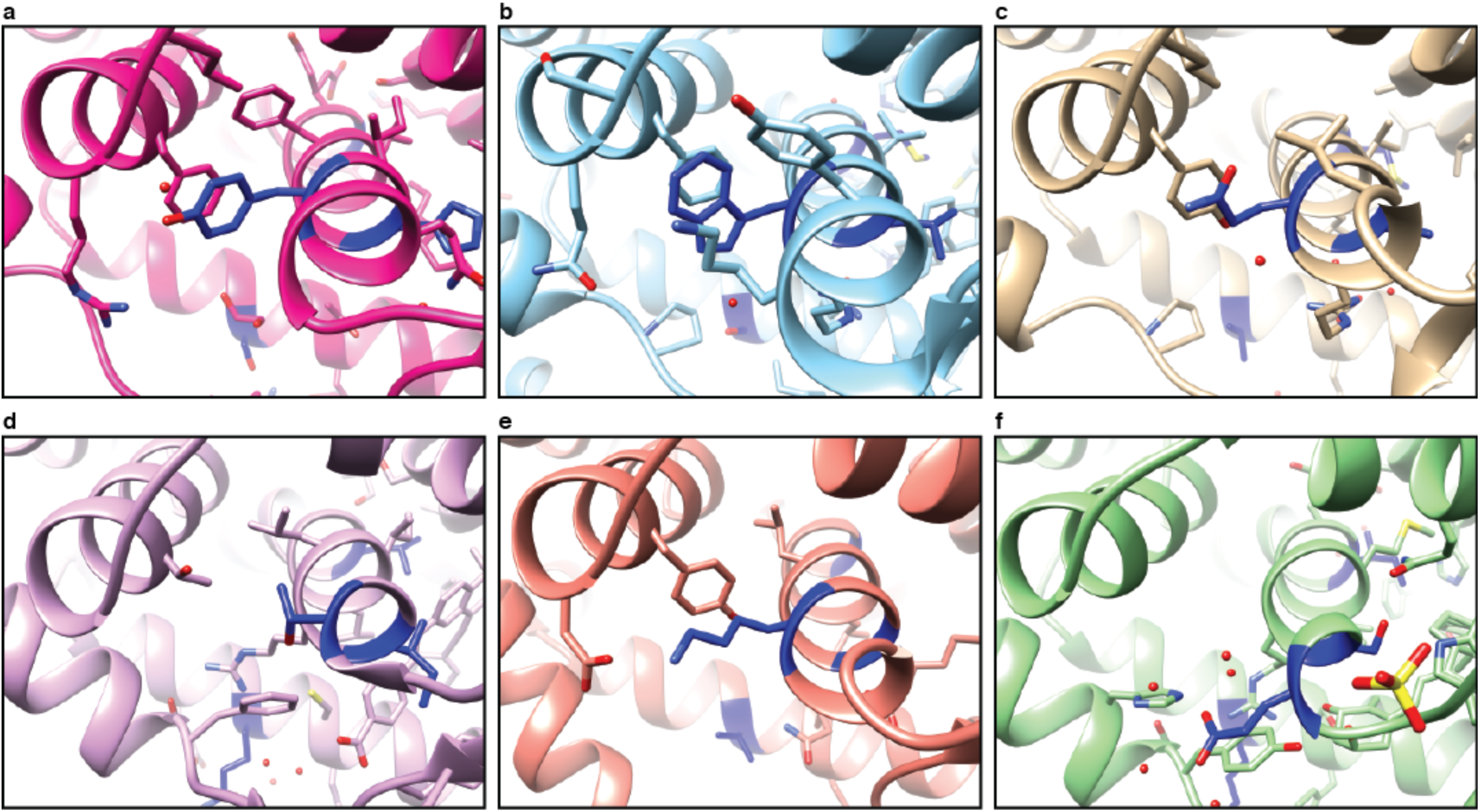
Local environments of the UMD residue of H5 distal to the ligand binding pocket (blue). π-π interactions between residues a. Tyr A312 and Phe A368 in SF-1 (4qk4),^42^ b. Trp A765 and Phe A818 of PR (2w8y)^43^ and c. Gln A371 and Tyr A422 of ERR*α* (1×b7).^44^ Equivalent residues also form salt-bridge interactions with H8 illustrated by e) Lys A185 and Asp A233 of HNF-4*γ* (1lv2).^45^ In other proteins these strong, specific interactions are replaced with general hydrophobic contacts such as in d. Thr B275, which is in contact with both Phe B199 and Thr B326 of RAR*α* (3kmz)^46^ and the same interactions are observed in RAR*γ* (e.g. see 1fcx, not shown). f. Lastly, the negatively charged Glu A277 found in this position of VDR (3ogt)^39^ forms a potential salt-bridge with His A139 and pi-pi interactions with Tyr A143. This results in a radically different interaction topology where the site binds to a different helix. Figure created with UCSF Chimera^36^ and Jalview.^35^

**Extended Data Table 1:** Differences in the structural properties of unconserved residues differentiated by their human missense variation classification.

| <u>Residue Counts</u><sup>a, e</sup> | Observed in<br>one or more mapped PDB |  | Not observed<br>in any mapped PDB |  | OR<sup>c</sup> | *p*<sup>c</sup> |
| --- | --- | --- | --- | --- | --- | --- |
|  | UMD<sup>d</sup> | UME<sup>d</sup> | UMD | UME |  |  |
| Ligand | 765 | 1,448 | 5,579 | 14,454 | 1.37 | $6.4 \times 10^{-11}$ |
| Domain | 649 | 1,312 | 5,695 | 14,590 | 1.27 | $3.5 \times 10^{-06}$ |
| Ligand, domain or nucleotides | 1,338 | 2,549 | 5,006 | 13,353 | 1.40 | 0 |
| Core | 1,635 | 1,995 | 4,709 | 13,907 | 2.42 | 0 |
| Part-exposed | 2,584 | 4,526 | 3,760 | 11,376 | 1.73 | 0 |
| Surface | 3,213 | 11,742 | 3,131 | 4,160 | 0.36 | 0 |
| <u>Column Counts</u><sup>b, e</sup> |  |  |  |  |  |  |
| Ligand | 156 | 407 | 114 | 357 | 1.20 | 0.23 |
| Domain (inter-chain) | 131 | 328 | 139 | 436 | 1.25 | 0.18 |
| Ligand, domain or nucleotides | 201 | 494 | 69 | 279 | 1.59 | 0.0033 |
| Core | 179 | 406 | 91 | 358 | 1.73 | $2.0 \times 10^{-04}$ |
| Part-exposed | 231 | 607 | 39 | 157 | 1.53 | 0.03 |
| Surface | 253 | 735 | 17 | 29 | 0.59 | 0.12 |

**Extended Data Table 2:** Example proteins with protein, ligand or nucleotide binding interactions involving residues in unconserved-missense depleted (UMD) columns from selected families (see Supplementary Data Table 1 for all families with discovered UMD columns).

| Family | Col.<sup>a</sup> | Res.<sup>b</sup> | Protein<sup>c</sup> | Ligand<sup>c</sup> | Nucleotide<sup>c</sup> |
| --- | --- | --- | --- | --- | --- |
| PF00001 | 525 | 89 |  | AA2AR_HUMAN (5iu8) [6] |  |
|  | 575 | 98 |  | ACM3_RAT (4u14) [6] |  |
|  | 584 | 99 |  |  |  |
|  | 780 | 129 |  | 5HT1B_HUMAN* (4iar) [23] |  |
|  | 792 | 130 |  | 5HT1B_HUMAN* (4iaq) [23] |  |
|  | 808 | 131 |  | 5HT2B_HUMAN (4nc3) [4] |  |
|  | 818 | 132 |  | 5HT2B_HUMAN (4nc3) [7] |  |
|  | 832 | 134 |  | ACM2_HUMAN (4mqs) [7] |  |
|  | 1075 | 176 |  | 5HT2B_HUMAN (4ib4) [8] |  |
|  | 1141 | 187 |  | AA2AR_HUMAN (2ydo) [4] |  |
|  | 1328 | 211 | ACM3_RAT (4u15) [2] | AA2AR_HUMAN (4eiy) [6] |  |
| PF00076 | 291 | 148 | B3GWA1_CAEEL* (5ca5) [13] | B3GWA1_CAEEL* (5ca5) [10] | CELF1_HUMAN (3nmr) [29] |
| PF00104 | 190 | 715 | ESR1_HUMAN (2jf9) [4] | A0A0B4J1T2_HUMAN* (2amb) [17] |  |
|  | 312 | 743 | NR4A1_HUMAN (3v3e) [3] | ANDR_HUMAN (1e3g) [35] |  |
|  | 330 | 750 | NR4A1_HUMAN (3v3e) [2] | ANDR_HUMAN (1t5z) [39] |  |
|  | 332 | 752 | NR1I3_MOUSE (1xnx) [3] | A0A0B4J1T2_HUMAN* (1t5z) [12] |  |

## Supplementary Data

**Supplementary Data Table 1:**
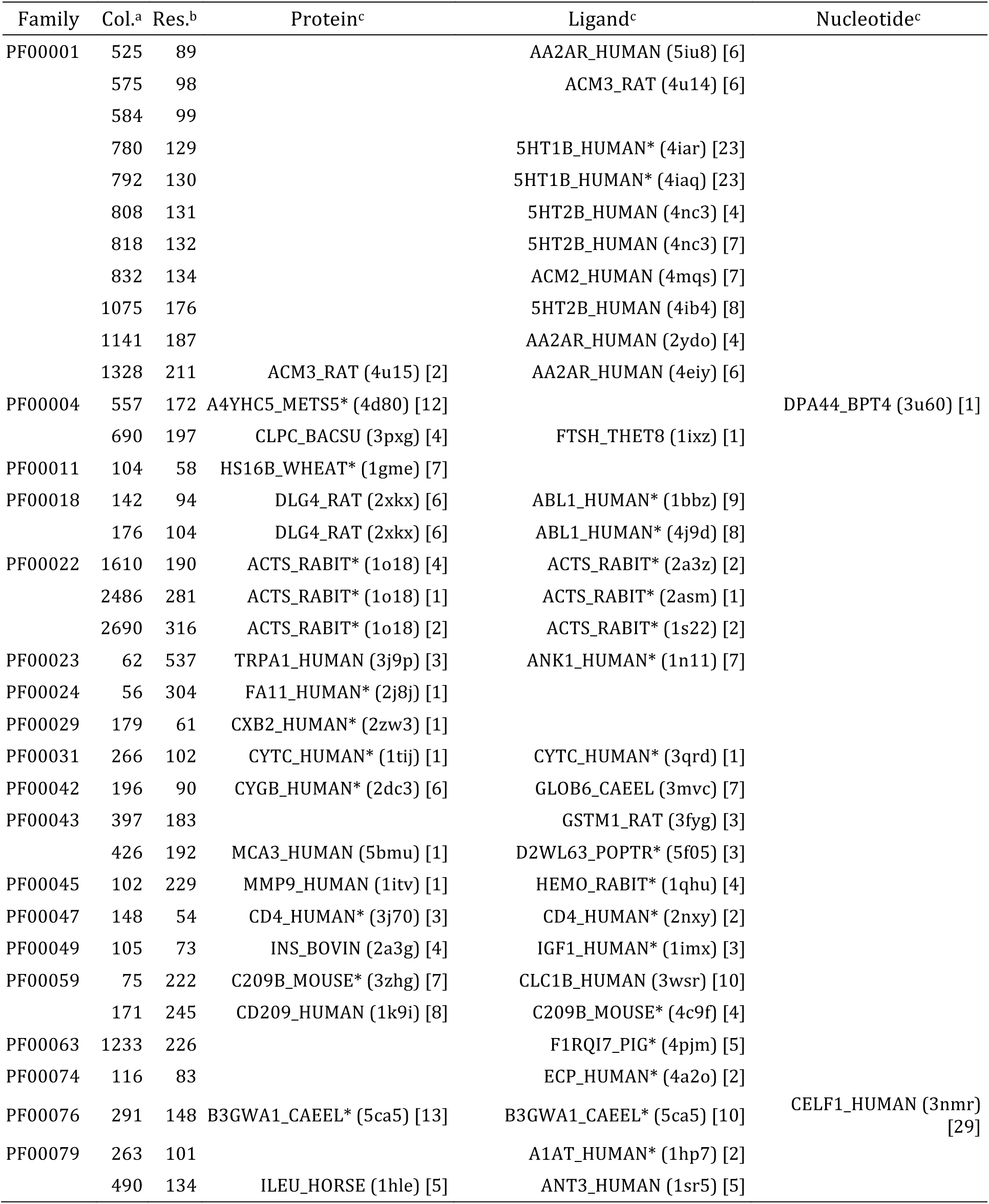
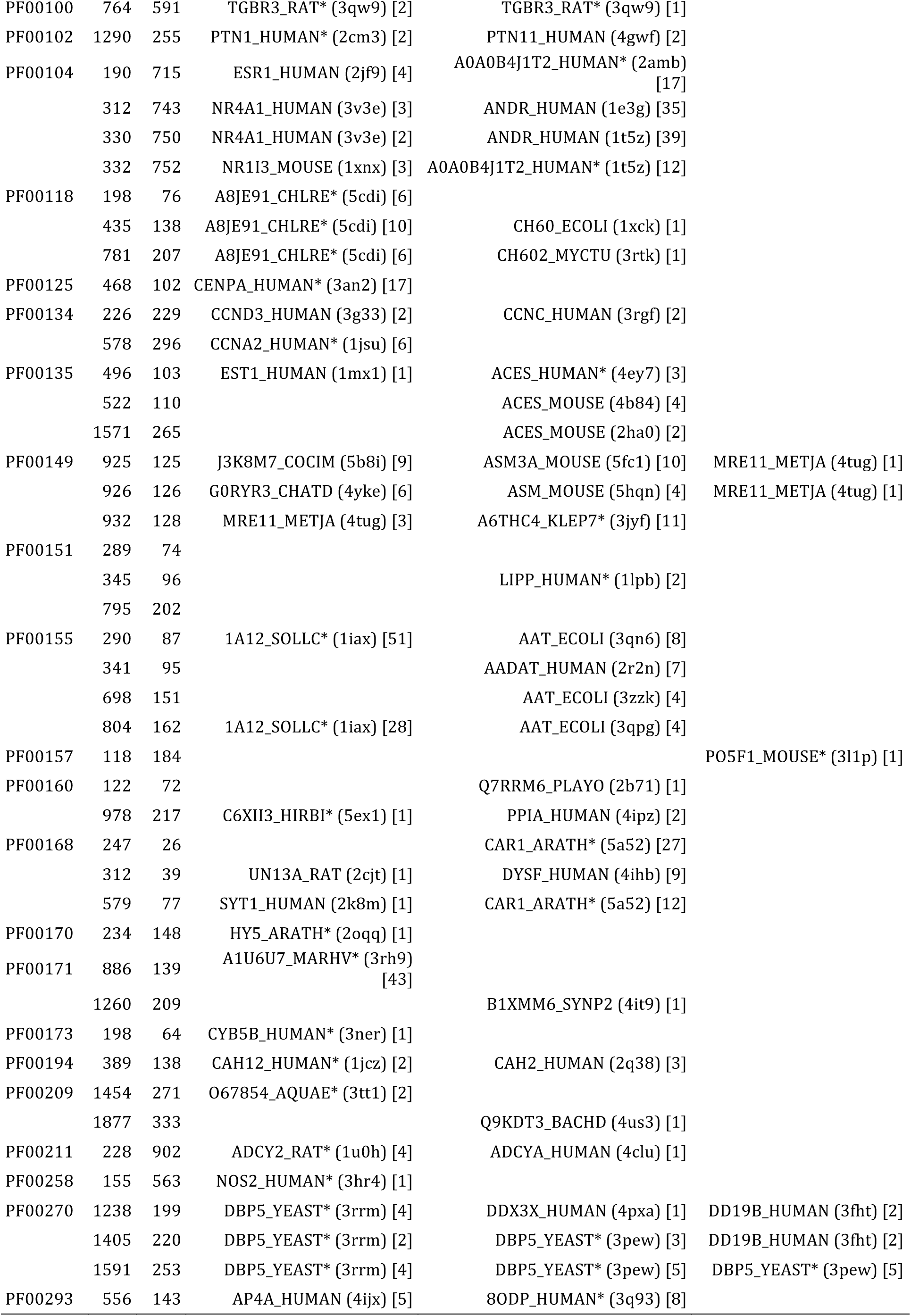
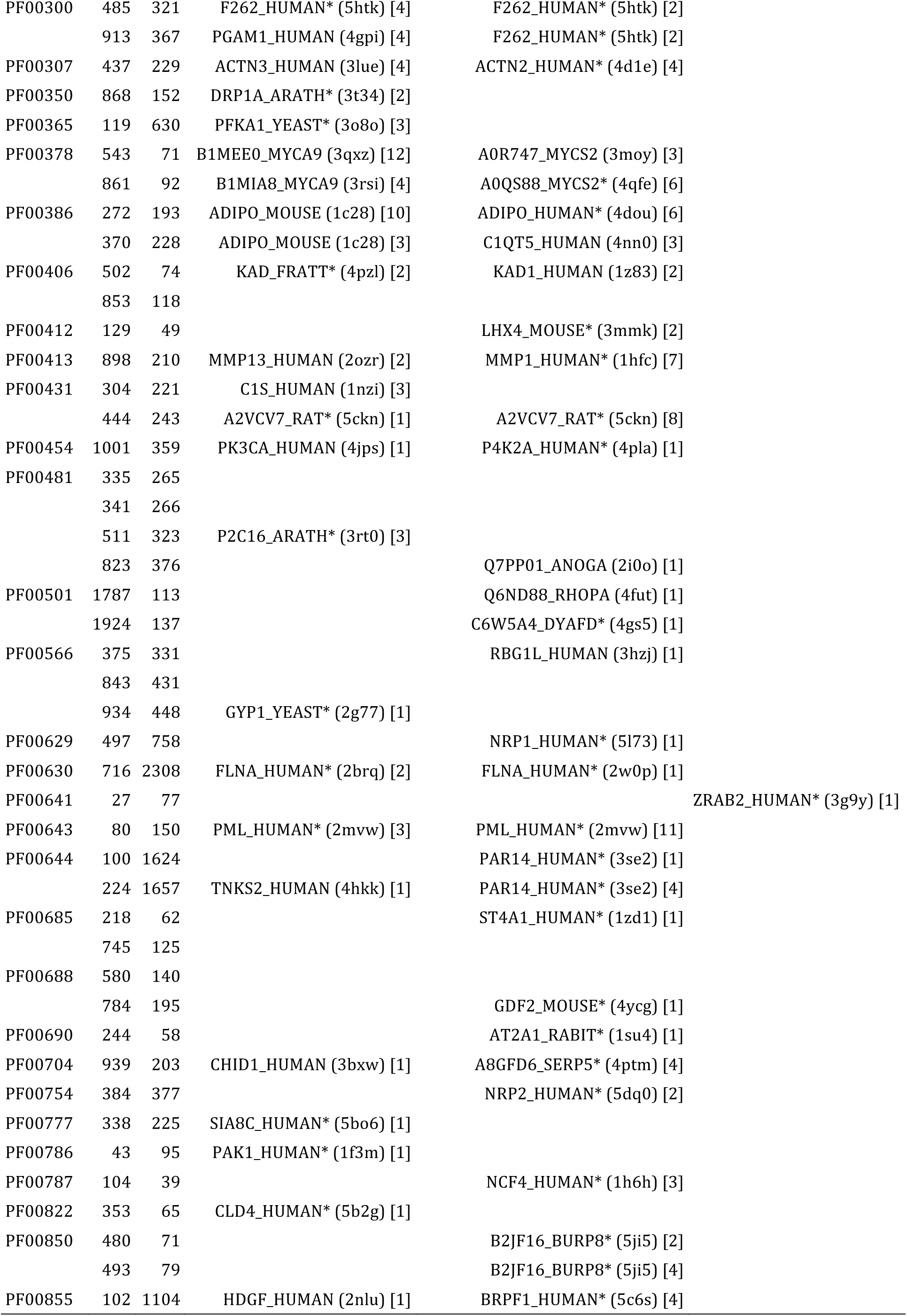
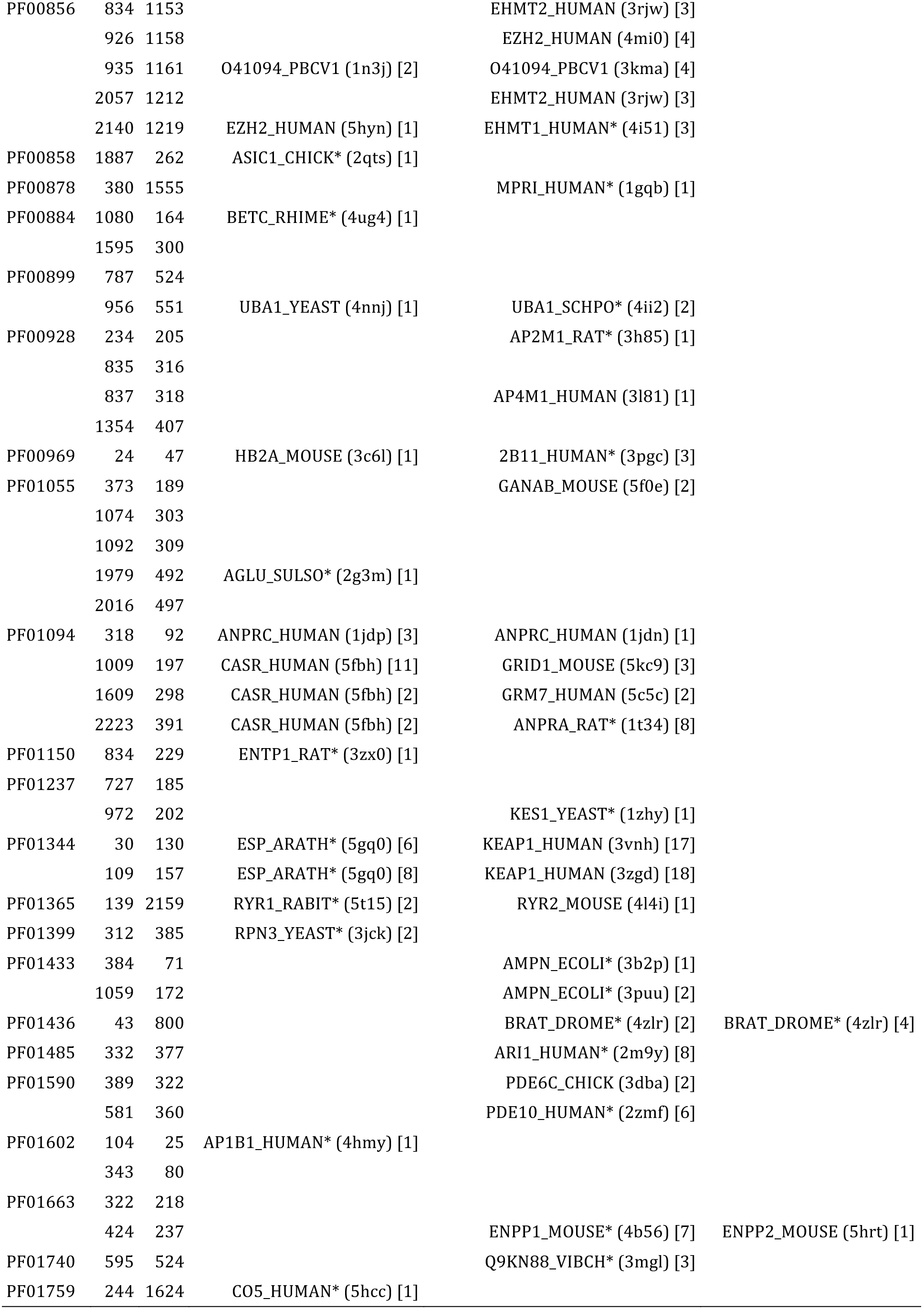
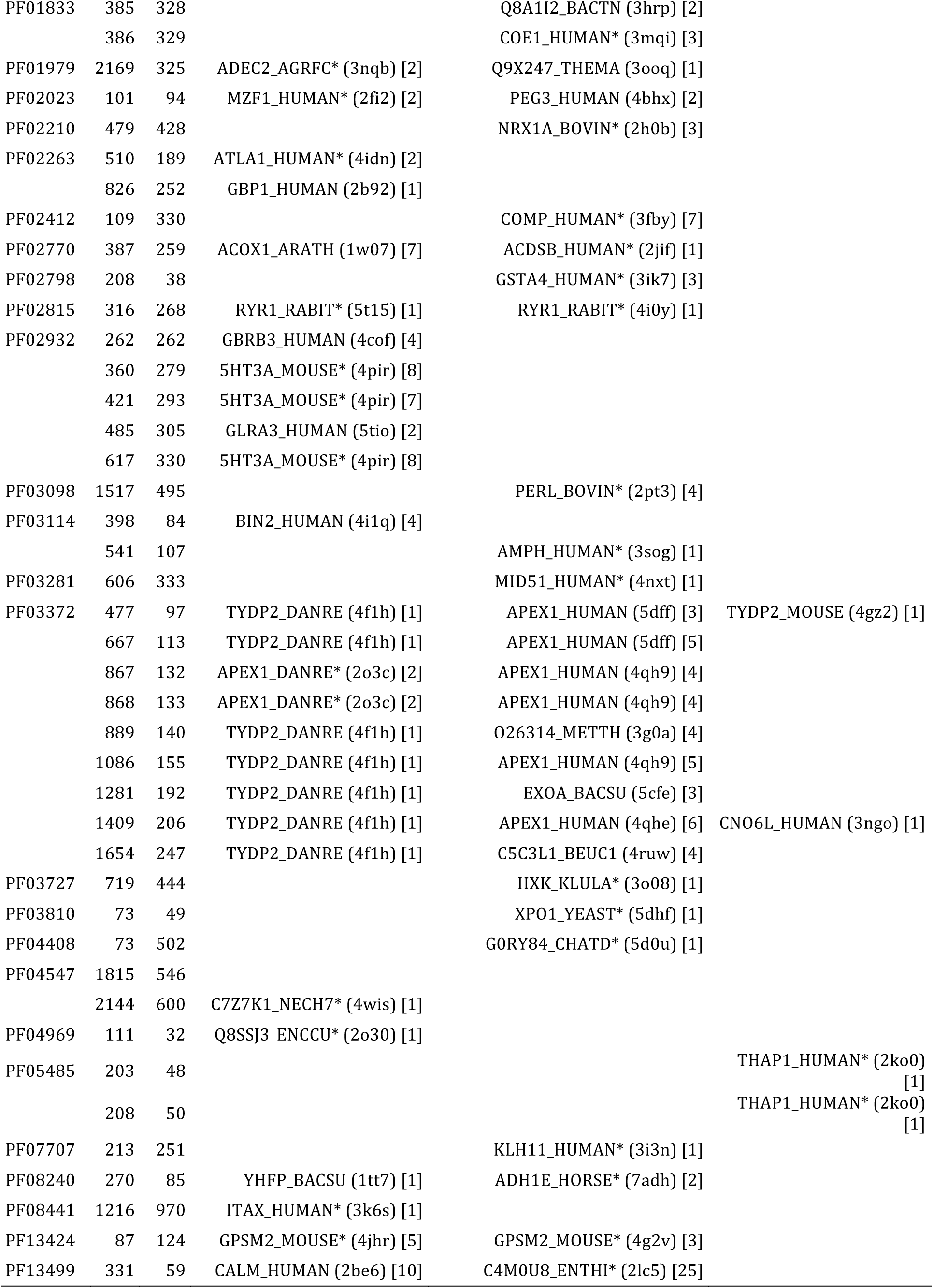
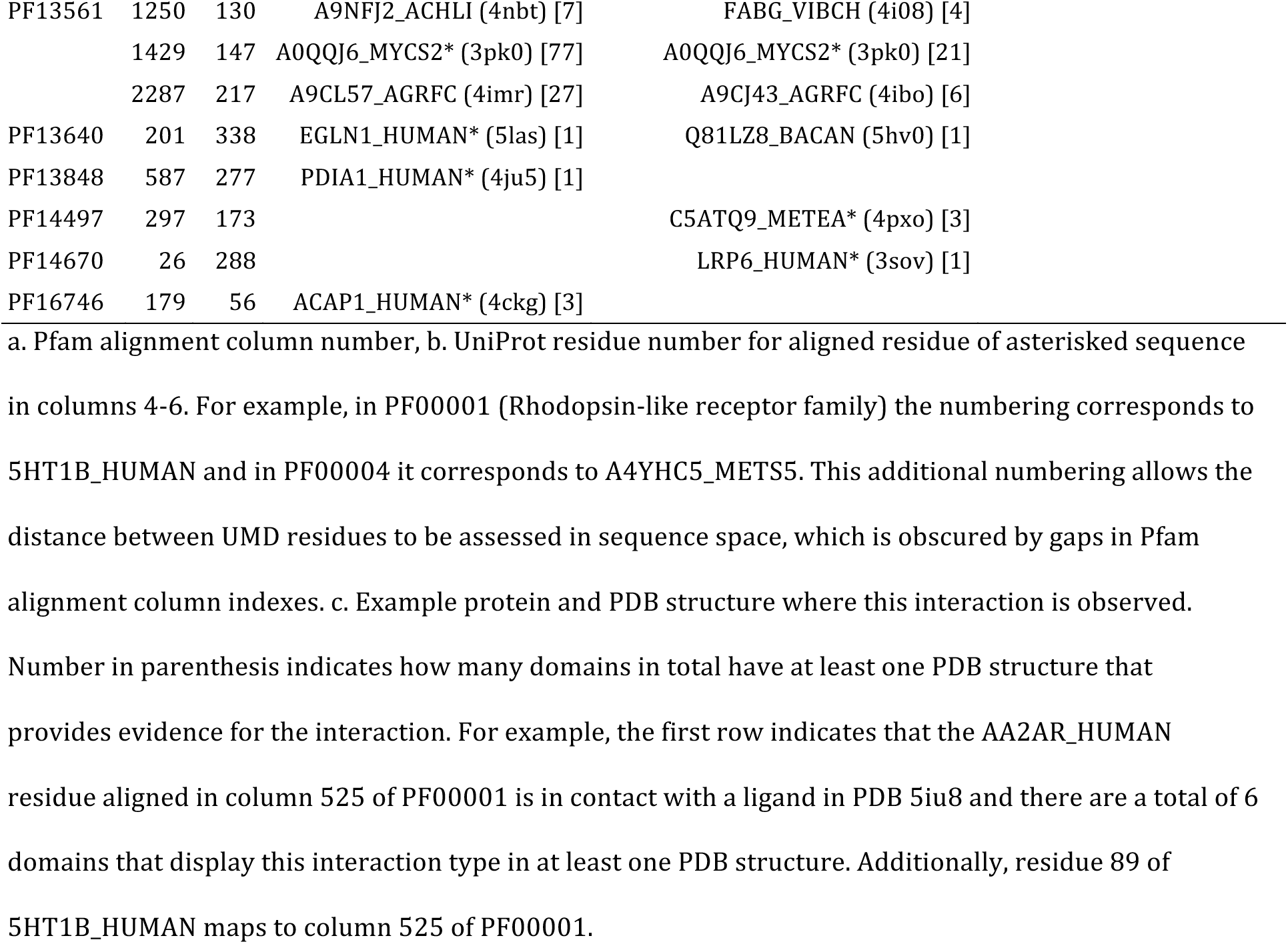
Example proteins with protein, ligand or nucleotide binding interactions involving residues in unconserved-missense depleted (UMD) columns. See table end for footnotes.

